# Phylogenetic analysis of molecular pathways in Tregs Suppressor Function

**DOI:** 10.1101/2021.07.30.453759

**Authors:** Michel-Edwar Mickael, Norwin Kubick, Pavel Klimovich, Irmina Bieńkowska, Suniti Bhaumik, Jarosław Olav Horbańczuk, Mariusz Sacharczuk, Rajatava Basu

**Affiliations:** Department of Experimental Genomics, Institute of Animal Biotechnology and Genetics, Polish Academy of Science, Postępu 36A, 05-552 Jastrzebiec, Poland; Department of Immunology, PM Forskningscentreum, 17854 Ekerö Stockholm, Sweden; Department of Biochemistry and Molecular Cell Biology (IBMZ), University Medical Center Hamburg-Eppendorf, Martinistraße 52, 20246 Hamburg, Germany; Bevill Biomedical Sciences Research Building, The University of Alabama at Birmingham, Birmingham, AL 35294-2170, USA; Department of Pharmacodynamics, Faculty of Pharmacy, Warsaw Medical University, l Banacha 1, 02-697 Warsaw, Poland

## Abstract

Treg-mediated suppression of conventional T cells is a fundamental step in regulating the adaptive immune response. It is known that Treg first appeared in vertebrates. However, little is known about the of major suppression pathways mediated by Tregs. We employed artificial intelligence text mining system to highlight the suppression pathways currently known to be utilized by Tregs. Our system identified various pathways mediated by CTLA4, calcium signaling, NfkB and NFAT. We simultaneously employed detailed phylogenetic analysis including multiple sequence alignment, phylogenetic tree building, ancestral sequence reconstruction, neutrality tests and positive selection test to investigate the evolution of Treg mediated pathways. We found that CTLA4 first appeared in vertebrates possibly arising from an IGV containing protein in cartilaginous fish. Conversely, we found that Tregs repurposed ancient pathways such as Calcineurin and CAMP Response Element Modulator. Interestingly, these two pathways were highly conserved between vertebrates and lower invertebrates indicating conservation of function. Taken together, our research indicate that Tregs developed its regulatory systems through evolution in vertebrates as well as reusing conserved ancient regulatory systems that are related to the innate immune system.

## Introduction

### Suppression of immune response important for addressing diseases pathology

Tregs plays a fundamental role in regulating the adaptive immune system [1] [2][3]. Tregs belong to CD4+ T cells family, that include Th1, Th2, Th17,Th9 and Th22. The importance of Tregs came to light, when it was discovered it plays a role in tolerance of transplanted organs[4]. Molecular investigations aiming to discoverer T reg differentiation and function increased dramatically following Tregs discovery. It was shown that TGFb plays an important in driving Treg phenotype from Th0. Afterwards the interconnection between Th17 and Tregs transcriptomic profile was highlighted. It was shown that both Tres and pathogenic Th17 are influenced by TgfB, smad pathway as well as other common molecular routes. However only Tregs as well as other intermediate cellular conditions such as Tre1were shown to have the ability to suppress other CD4+ T cells also known as conventional T cells.

### Treg suppression mechanisms

Tregs suppression mechanisms are widely investigated, there are various indirect mechanisms with which Tregs suppress T con through indirect pathways including CTLA4, where it was shown that CTLA4 compete with CD28 on the interaction with CD80/CD86. Other indirect pathways include CD40/CD40L, neuropilin-1 and A20. Although Tregs are also known to use various cytokine to suppress other T co including TGFb, IL10,IL27 and IL2. Another mechanism is done through manipulating other main T con main transcription factors such as RORgt in Th17. However, we and others have found that RORgt play a role in sustaining FOXP3 suppressive functions. Furthermore, it was shown that Tregs uses ICER to suppress NFAT pathway. Also Tregs have been shown to sown regulate calcium pathway needed for conventional T cells differentiation through a Calcineurin (PPP3CA) dependent pathways.

### Treg evolution not fully understood

From an evolutionary point of view, Tregs suppression mechanisms are still not understood. It is well known that Tregs, as well as other T cells first, appeared in vertebrates. However, it is still not known if the pathways used by Tregs are its innovation, or where they were acquired from previous organisms. It is also known that adaptive immunity seems to have first in vertebrates. However, the evolution mechanism of the regulation adopted by the adaptive immune system has not been yet understood. Furthermore, the origin of these pathways is still not known. There is strong debate about the origin of regulation in vertebrates, One side of the argument suggests that regulatory mechanisms are universal, and they could have evolved from one initial mechanism. The other side of the argument suggests that the evolution of regulation of cells is divergent with multiple local mechanisms appearing in various vertebrates. Furthermore how lampreys mediated control over their immune-like cells is still unknown. Lampreys are known to posse two types of immune-like cells that are affected by cytokines, exist in the blood, and have pro and anti-effects. However, with mechanisms existing behind their regulation is still unknown. Taken together these observations suggest there is ne a need for better understanding of the evolution of the adaptive immune system regulations, materialization of Tregs suppression mechanisms.

### What we have done

We investigated the evolution of Tregs mediated pathways through performing phylogenetic analysis. We downloaded proteins sequences relates to Tregs functions from humans GEO PubMed proetin repository. Using Blastp we identified homologs for these sequence in 12 species and orders. After that we reconstructed ancestor sequences for known Tregs suppressors and investigated the evolutionary selection of these suppressors using PAML. Our results show that Tregs have adopted both ancient and novel mechanisms to acquire the ability to suppress T cons. Furthermore, our results support the argument that the origin of regulation in the adaptive immune system did not first appear in vertebrates. However regulation in the adaptive system is not unique and it appears through different divergent mechanisms starting from early vertebrates. Finally we found several suppressors mechanisms in lampreys, which suggest that lampreys possess a Treg like cells suppression mechanism.

## Methods

### 2.2. Database Search

The focus of this research was investigating the IL family’s evolution and their origins. Due to the diverse nature and long evolutionary history of the IL families, we used protein sequence alignment. Moreover, to make sure that our analysis is a reasonable representation of IL evolutionary history, we investigated the presence of each of the family members using rodents, Monotremes, Marsupialia, Diapsida, Actinopterygii, Salientia, Caudata, Petromyzontiformes Tunicates,Arthropods, Nematoda, Spiralia, Cnidaria and Placozoa, which span more than 500 million years. Human protein identified by our AIST system were used for BLASTP searches against the above-mentioned proteomes. The longest transcript was used in the analysis. Sequences were selected as candidate proteins if their E values were ≤1e−10. Sequences were further filtered by comparing the conserved domain in each protein investigated [5] [6] [7].

### 2.3. Alignment and Phylogenetic Analysis

Phylogenetic investigation was done in two steps. First, IL family amino acid sequences were aligned using MAFFT using the iterative refinement method (FFT-NS-i). After that, we employed, PHYML implemented in Seaview with five random starting trees to generate the final tree. The Neutrality test was performed using MEGA 6.

### 2.4. Ancestral Sequence Reconstruction (ASR)

We applied the maximum likelihood method to infer the ancestral sequence of each of the proteins investigated. For each protein, we used the ASR algorithm implemented in MEGA6 to build ancestral sequences This was followed by BlastP against the nearest earlier diverging organism. The BlastP outcome was only accepted if the E-value threshold was less than e−10. The evolutionary network for ancestral sequences was built using SplitsTree with a default setting and bootstrap value of 100[8].

### 2.5. HHsearch

The HHsearch method was used to examine the evolutionary history of the ILs. Only proteins that have already diverged before the most ancient members of the family were considered as candidate parents [2,27].

### 2.5. Positive Selection

We used the maximum likelihood algorithm in PAML to identify ILs that have gone through positive selection[9]. In the first instance, respective complementary DNAs (cDNAs) were estimated using the backtranslation function on the EMBOSS server (https://www.ebi.ac.uk/Tools/st/emboss_backtranseq/ (accessed on 5 May 2021) [32]. Next, we employed the CODEML PAML v4.4 program to evaluate global and branch selection by calculating the substitution rate ratio (ω) given by the ratio of nonsynonymous (dN) to synonymous (dS) mutations[10][11].

## Results

| Pathway | Genes |
| --- | --- |
| **CTLA4** | CD28,CD80,CD86 and LFAI-1 |
| **Indirect pathway** | A20, CD40-CD40, neuropilin-1 |
| **Suppressive cytokines** | TGFb, IL10,IL35 |
| **Induce apoptosis** | IL2, TRAIL, CD3,CD46,CD25,bim, Puma |
| **Upregulation of TF** |  |
| **Adenosine** |  |
| **cAMP, ICER, and NFAT** |  |
| **TCR** |  |
| **Ap1** |  |
| **NFAT** |  |
| **NFKB** |  |
| **Calcium signaling** |  |

### Diverse suppressor mechanism evolved in Tregs

#### CTLA4 perform its function through four different pathways

(i) CTLA4 expressed on Tregs was shown to compete with CD28 expressed on T con cells to limit activation of conventional T cells through binding between CD80/CD86 on APC and CD28 on T con cells through a process called trans-endocytosis [12]. (ii) It has been claimed that this line of action is dependent on LFA1-ICAM1 interaction [13]. (iii) Also it was shown that CTLA4 could increase the expression of IDO in APC which causes T con starvation. (iv) Additionally CTLA4 can also reduce the expression of GCLC and GSS which are responsible for glutathione synthesis leading to forming a redox environment that is unfavorable for T con growth [14]. Interestingly CTLA4 was not expressed beyond vertebrates, same as CD80 and CD28. However, CTAL4 is expressed in the earliest lung fish related to tetrapoda (e.g. *Latimeria chalumnae* (Coelacanth), accession number AFYH01096763.1). Additionally, we found three homologs of CD86 in athropoda, namely Ostrinia furnacalis, Vanessa tameamea] and [Pieris rapae] (XP_028167410.1, XP_026493316.1 and XP_022129956.1) (figure 1). LFA1 first appeared in Tunicates. Interestingly, we found that IDO as well as GSS and GCLC first appeared in Profieria (sponges) (figure 1).

**Figure 1.**
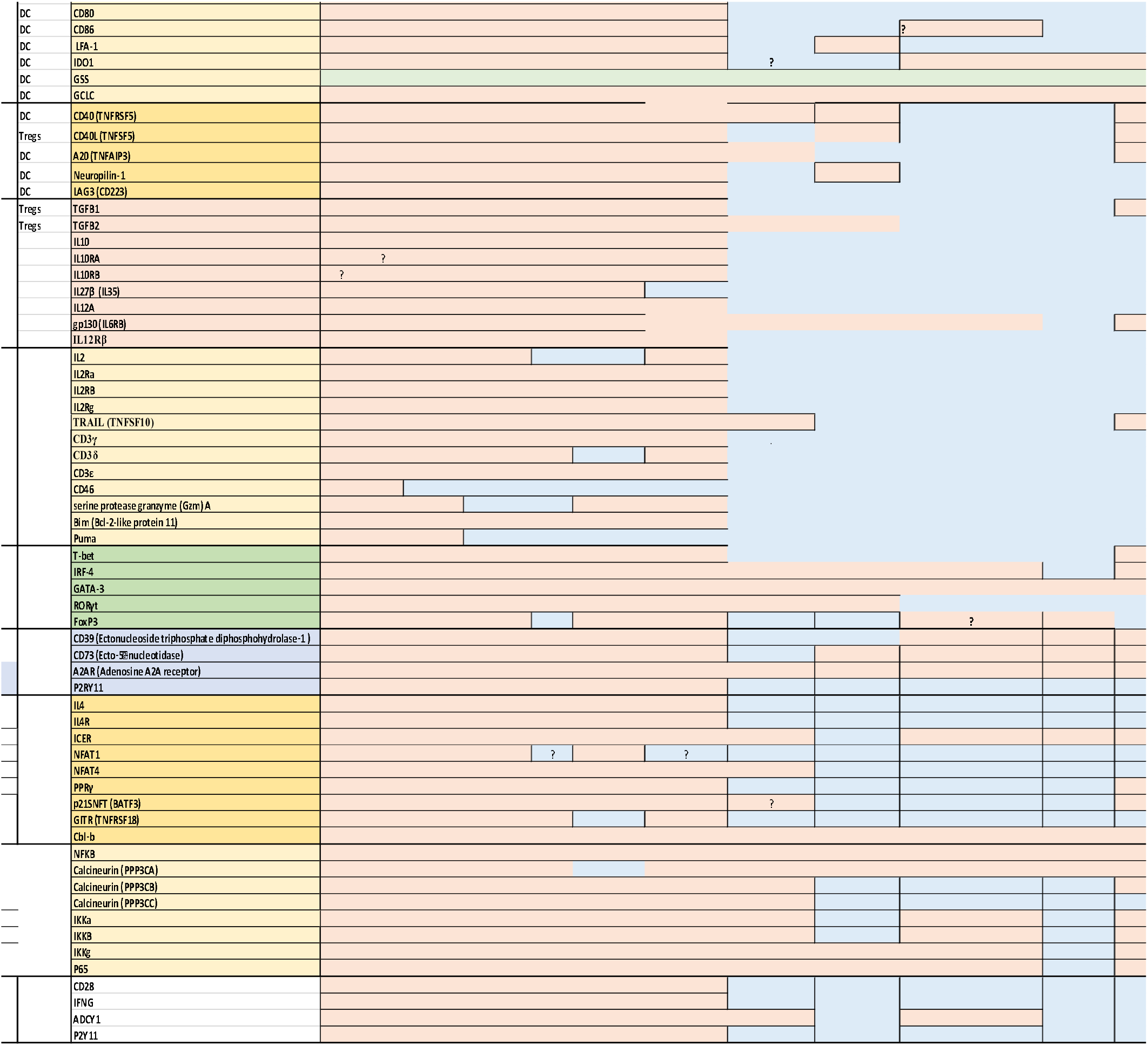
Evolutionary history of Treg mediated suppression pathways.

**Figure 2.**
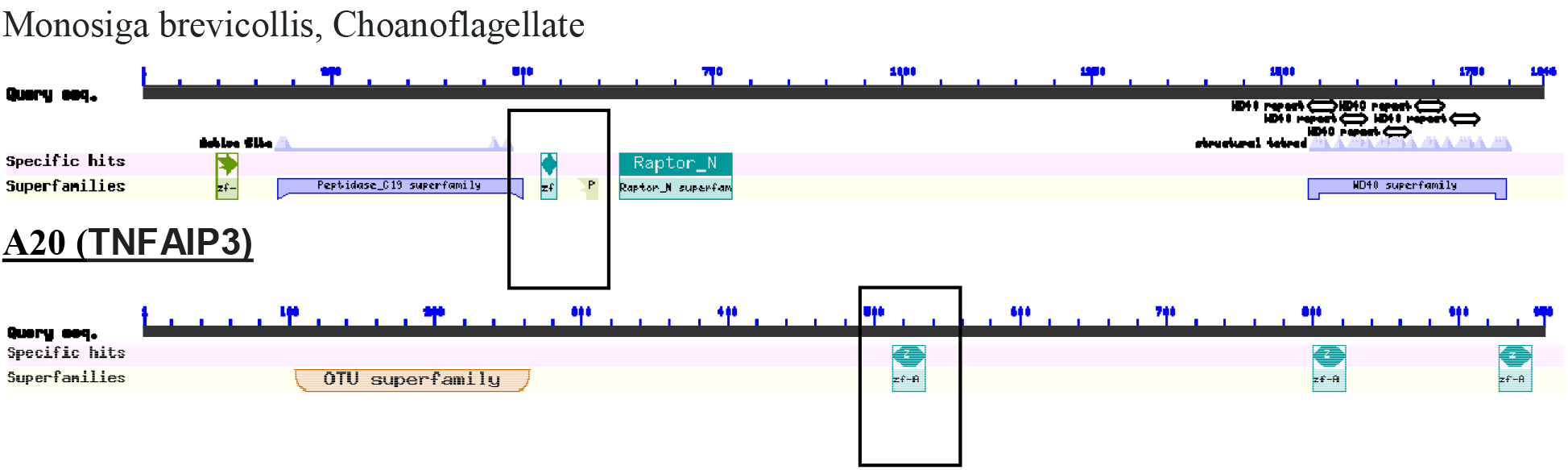
Origin of A20 gene. A20 origin shares a A20-zinc finger like with the Choanoflagellate *Monosiga brevicollis*

Another pathway of indirect pathway include the interaction between CD40L expressed on Tregs and CD40 expressed on DC that inhibit the function of the DC [15]. Interestingly, we found that CD40 and CD40L first existed in Mollusca (figure 1). Another molecule that also used by Treg to inhibit DC is A20, also its role is still not clear. A20 is a Ubiquitin-editing enzyme that contains both ubiquitin ligase and deubiquitinase activities and is an essential protein in the Ubiquitin complex. It promotes the termination of the TNF-mediated activation of NF-kappa We found that A20 is an ancient proetin as it has appeared during the emergence of sponge, while neuropilin-1 first appeared in Cnidaria (figure 1). Conversely LAG3 which was shown to be expressed on Tregs and binds to MHCII on denderditc cells [16], seems to have diverged during the emergence of fish.

#### Tregs suppressive cytokines include four

cytokiens mainly TGFβ (including TGFβ1 and TGFβ2) as well as IL35 and IL10 (figure1). Our results indciate that TGF*β* homologs are ancient. TGFβ1 first appeared in sponge, while TGFβ2 first appeared in Cnidaria. IL10 and IL35 in agreemnt with [10] we found that IL10 as well as IL10Ra and IL10Rb, diverged during the emergence of Actinoptyregii. IL35 is composed of IL-27β and IL-12α, its receptors are gp130 and IL-12Rb GP130 first appeared in Mollusca while IL27β, IL12A and IL12Rβ diverged in fish (figure 1) and [10].

#### Treg mediated apoptosis induction pathways

include IL2 and its three receptors IL2Rα, IL2Rβ and IL2Rγ first diverged in fish. We were also able to locate TRAIL gene in lampreys as well as invertebrates (including Mollusca and cnidarian) (figure 1). CD3γ, CD3δ and CD3ε are the three protein forming CD3 part of the TCR complex. According to our results they seem to have evolved during the emergence of fish. Conversely CD46 which was shown to enhance Treg activation only appeared in rodents (figure 1). Once activated Tregs were reported to induce apoptosis in T con through GZMA mediated pathway [17], we found that GZMA has diverged in fish (figure 1). Additionally we found that Bim (Bcl-2-like protein 11) and Puma that were also implicated in Treg induction of apoptosis diverged during the emergence of fish and Marsupialia respectively (figure 1).

#### Treg mediated an indirect change in the main transcription factors of T con different populations

T con main population include Th1 and it is mediated by Tbet. We found that Tbet first diverged during emergence of Mollusca. IRF-4 was shown to be critical for the function of Th2 and Th17 and we observed that it first appeared in cnidarian. Gata3 appears in all investigated starting from cnidaria. RoRgt first appeared in tunicate, while FoxP3 which is the main TF of Tregs and it first diverged during the emergence of nematode.

Another way Tregs have been shown to suppress T con through hydrolysis of extracellular ATP to ADP or AMP through the enzyme CD39. We found that CD39 diverged during the emergence of placozoa. CD73 was also shown to inhibit T con through degrading AMP to adenosine. We identified CD73 homologs in all genus investigated from Sponges to humans except lampreys (figure 1). Interestingly, adenosine receptors seems to have diverged during Cnidarians emergence, while Purinergic Receptor are absent in vertebrates [18].

When Tregs suppress T con through augmenting cAMP. cAMP induces early repressor (ICER) to suppress IL4. We found that ICER was first expressed in Cnidarian (figure 1), while IL4 first appeared during the emergence of the Actinopterygii similarly, IL4RA diverged in Actinopterygii in agreement with [10]. Another pathway that is related to ICER is through NFAT which was shown to forms inhibitory complexes on cytokine promoters with transcriptional repressors such as ICER, PPRγ, or p21SNFT We were able to identify, NFAT4 in sponges, while NFA1 seems to be have recent diverged during emergence of **<u>xenopus/bony fish</u>** PPRγ, p21SNFT were more ancient and they both first appeared in Spiralia (figure1). Interestingly, as ICER induction in suppressed T cells was CTLA-4/B7 and GITR-dependent. GITR first appeared in fish (figure 1). Also, Cbl-b deficient T cells are less sensitive to suppression by Tregs. CBL-B was identified in all species from sponges to humans.

Another important aspects of Tregs mediated regulation of conventional T cells is suppression of calcium signaling. Calcineurin was suggested to inhibit NfkB, through a IKK mediated pathway.. Calcineurin has three family members, namely PPP3CA, PPP3CB and PPP3CC. Interestingly, whereas NFKB is as ancient as sponges, Calcineurin is more diverse with PPP3CA, PPP3CB emerging during sponges, while PPP3CC first appearing in Mollusca. IKK is also ancient, it has three members, which all emerged during invertebrates divergence (figure 1).

### Families of the Tregs mediated suppressors

Investigating the structure of Tregs mediated suppressors revealed that Tregs employs a heterogonous number of molecules as suppressors, these suppressors belong to various families (i) immunoglobulin super family and these include CTLA4 and LAG3.(ii) TNF super families, it associates and their receptors and it includes CD40L(TNFSF5), CD40, A20 (TNFAIP3), TRAIL (TNFSF10) and GITR (iii) neuropilins (iv) TGFB superfamily (v) Grnazymes (vi) E-NTPDase family of ectonucleotidases: (vii) Ecto-5’-nucleotidase (viii) bZIP transcription factor domain containing protein (ix) NFATs (x) 5-hydroxyicosatetraenoicacid and 5-oxo-eicosatetraenoic acid family (xi) Basic leucine zipper transcriptional factor ATF-like (xii) ubiquitin ligase and (xiii) Calcineurins family.

### Origin of Tregs mediated suppressors

The origin of Tregs mediated repressors pathways is diverse. Irrespective of its name CTLA4 is not related to CTLA1 (GRANZYME B), CTLA2 or CTLA3 (Granzyme A). Alternatively, it is nearest homolog is CD28. CTLA4 evolved from a protein containing IGV domain (99.5 5 and 1e0-17) (Table 2). LAG3 emerged from an immuunoglobin proetin (99%). The TNFα superfamily contains 19 members that bind to 29 members of TNF receptor superfamily [19]. Among its members that are known to be employed by Tregs as suppressors are CD40, TRAIL. tumor necrosis factor ligand superfamily member 10-like Tax=Lingula unguis 98.92. TNFa receptor family contains 29 receptors. We identified one homolog of TNFa in Monosiga brevicollis (Choanoflagellata) e-value 2e-6. The TNFAIPs family mainly include TNFAIP1, TNFAIP2, TNFAIP3, TNFAIP4, TNFAIP5, TNFAIP6, TNFAIP8 and TNFAIP9. (Blast value of 7e-04). Neuropilin seems to have diverged from a cub containing domains or a discoidin I. Granzymes are a large family consisting of 11 members, interestingly, only four granzymes appear in human genomes (e.g, A, B, K and M) [20]. Grnazymes could have diverged from Trypsin (1e-38), melanization protease 1 2e-33, or corin (3e-32). CD73 belong to 5’nucleotidases. Seven memebrs of the 5’-nucleotidases have been characterized (e.g., NT5C1A, NT5C1B, NT5C2, NT5C3, NT5C3L, NT5E and NT5M. They have been reported to be as ancient as bacteria [21][22]. We reconstructed the ancestral sequence for 5’-nucleotidases that could have been present in LUCA. CD39 belong to the E-NTPDase family (e.g., ectonucleotidases). Ectonucleotidase have eight members (NTPDase1, 2, 3, 4,5,6,7 and 8)[23]. We locate the origin of Ectonucleotidase to have exist in algae (Tbale 2). BATF3 (P21) belongs to the P21 family, which in turn consist of two more members namely BATF1 and BATF2. The coinstructed origin for the BATF, nearest homolog is gene jun dimerization protein 2-like that first appeared in Cnidaria [Actinia tenebrosa] (e-value 4e-06). There are five different NFAT members NFATc1, NFATc2, NFATc3, NFATc4, and NFAT5[24]. The origin of the NFAT seems to be similar to NFAT4C genes that first appeared ins sponges (Table 1). CBL family consists of three members namely C-CBL, CBL-B AND CBL-C. The nearest homolog to this family is E3 ubiquitin ligase Cbl TKB [Salpingoeca rosetta] (2e-101). In vertebrates, the gene family of PPAR consisted of PPAR*α*, PPAR*β* (also called PPARb/d or PPAR*δ*), and PPAR*γ* [25].

**Table 2.**
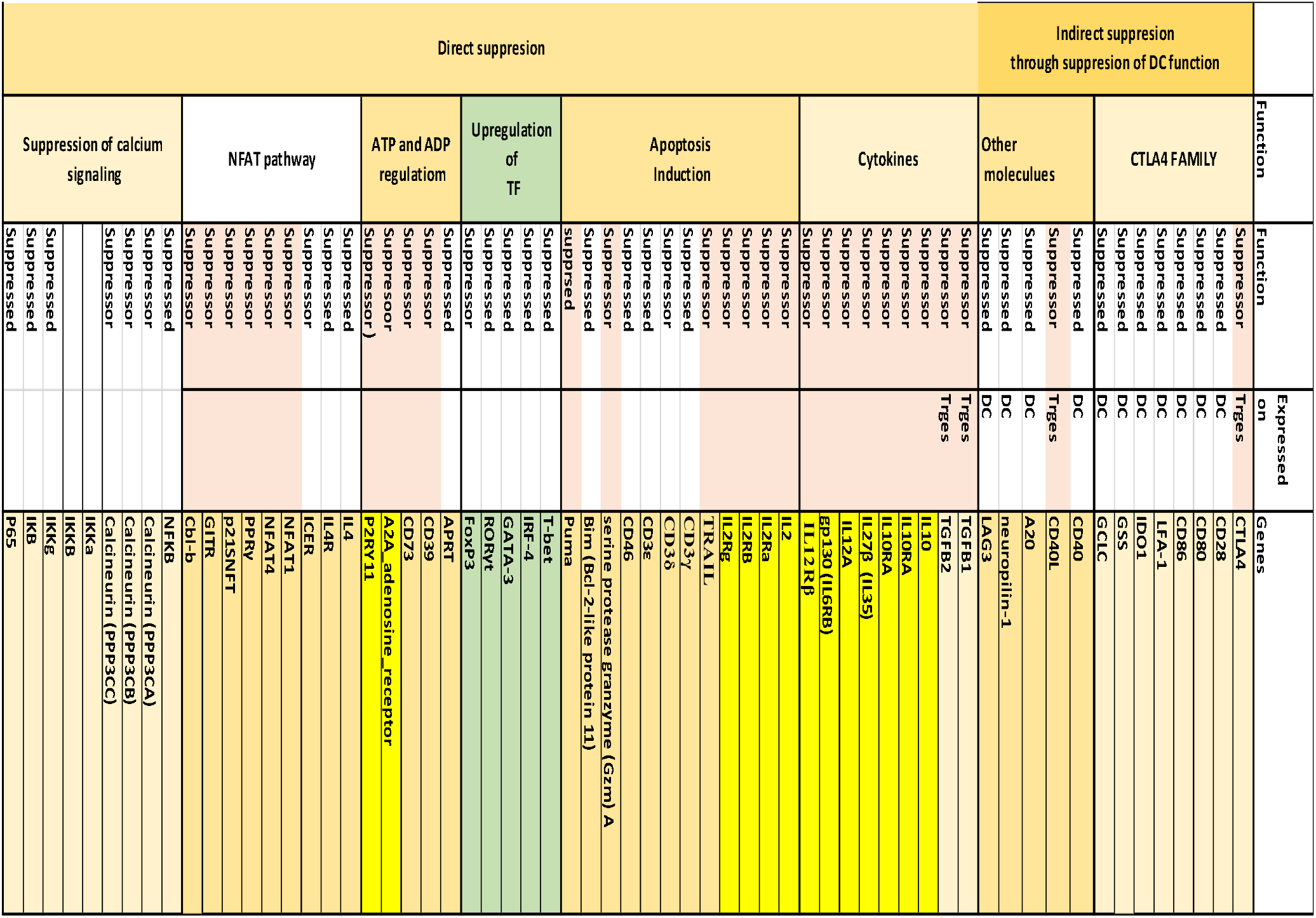
Treg suppression mediated genes categorization by function and expression.

**Table 3.**
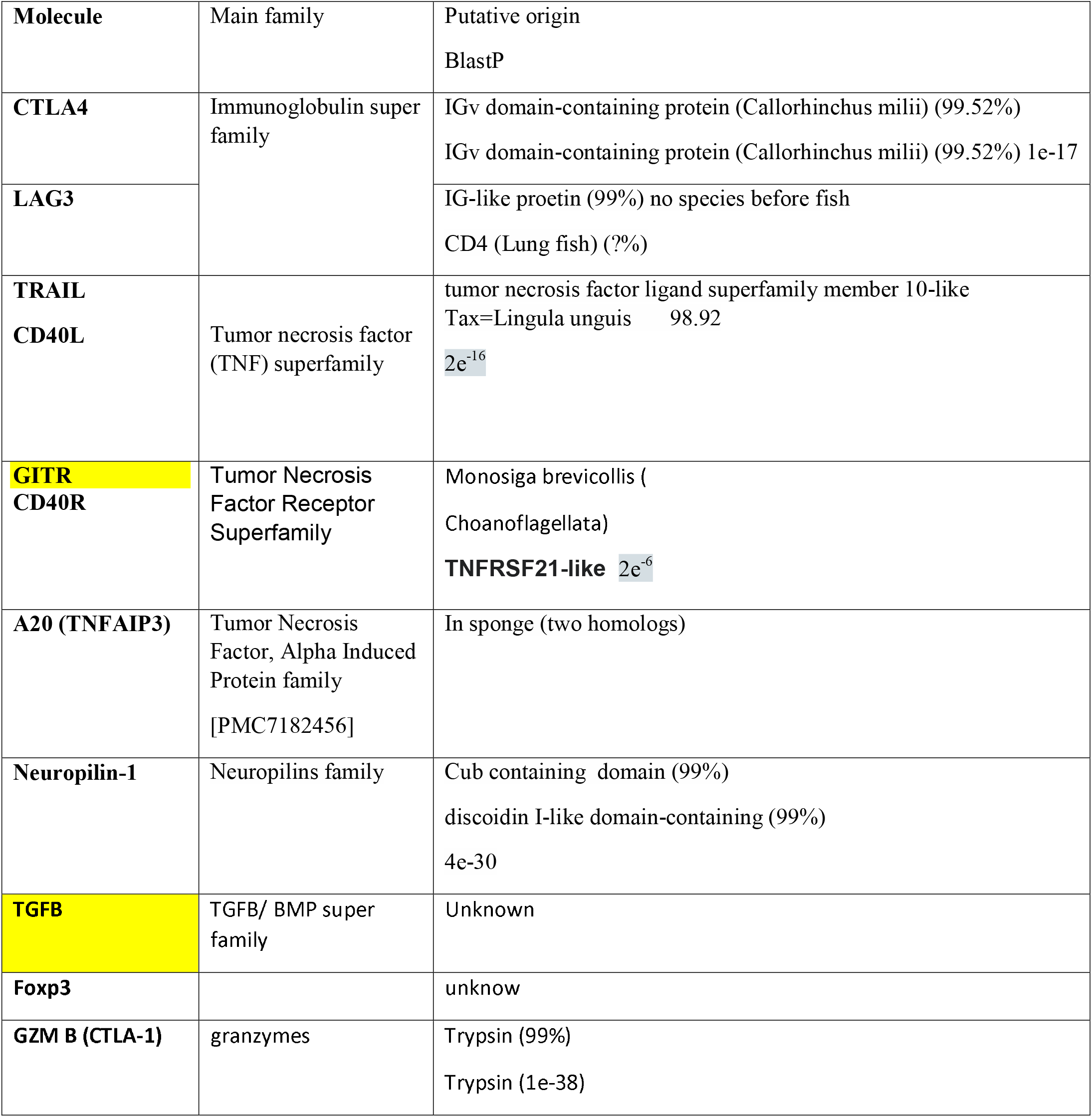
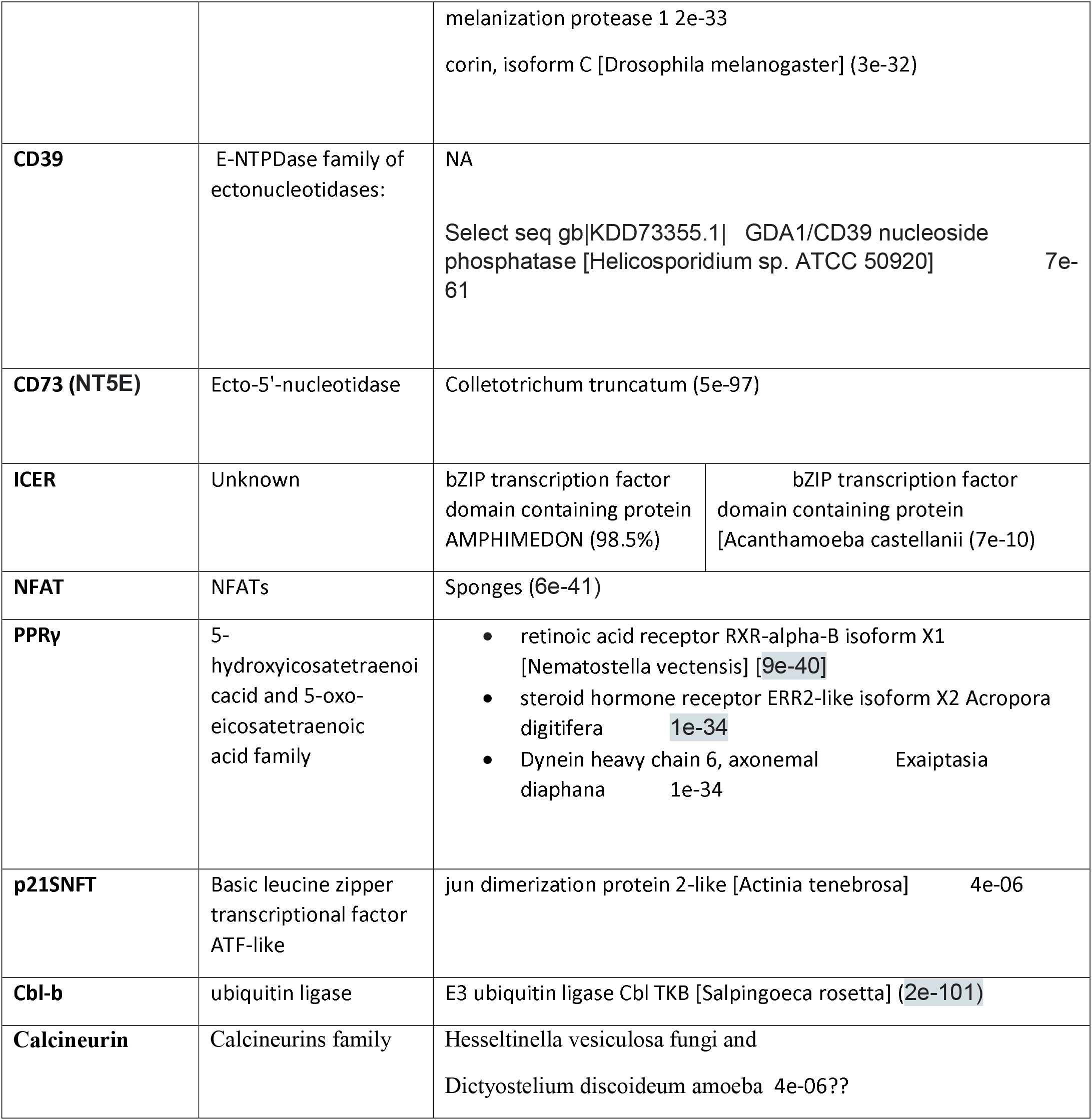
Putative origins for Tregs mediated pathways.

### Positive selection

We investigated positive selection among Treg mediated suppression pathways. We employed PAML for computation of global w value. Our results show that, there are various suppressions employed by Tregs that evolved under strict conservation mechanisms including CLAT4 (0.42, P-value <0.0005) as well as the CD40L and its receptors, in addition to TRAIL, GZM, CD73 and GITR. On the other hand several suppressors employed by Tregs were subjected to strong positive selection including TGFB (1.46, p-value < .00001), in addition to LAG3,CD39, NFAT and CBLB.

| Molecular | Global | P-value |
| --- | --- | --- |
| CTLA4 | 0.42 | <0.0005 |
| CD40 | 0.27 | <0.00001 |
| CD40L | 0.39 | <0.00002 |
| A20 (TNFAIP3) | 0.96 | not significant at $p < .05$. |
| Neuropilin-1 | 1.13 | not significant at $p < .05$. |
| LAG3 | 1.50 | < .00001 |
| TGFB | 1.46 | < .00001. |
| TRAIL | 0.44 | <.00006 |
| GZM | 0.31 | < .00001 |
| CD39 | 1.45 | < .00001 |
| CD73 | 0.48 | < .00001 |
| ICER | 0.63 | < .01 |
| NFAT | 1.88 | < .00001 |
| PPRγ | 0.61 | *not significant at p < .05.* |
| p21SNFT | 0.66 | *not significant at p < .05.* |
| GITR | 0.50 | < .0001 |
| Cbl-b | 1.38 | < .00001 |
| Calcineurin |  |  |

## Discussion

### Tregs mediated mechanisms evolution are diverse

The evolution of Tregs mediated suppressors is diverse and they were adopted from various ancient as well as novel pathways. Tregs seems to have used three strategies in acquiring its suppression ability (i) adapted ancient suppression pathways to suppress ancient and novel T con pathways and acquired novel pathways to suppress novel and ancient pathways. Our algorithm was able to identify various pathways with which Tregs suppress conventional T cells (i) indirect through the CTLA4 pathways (ii) in directly through other molecules (iii) using cytokines (iv) inducing apoptosis (v) upregulation of TF (vi) NFAT pathway and (vii) Suppression of calcium signaling. CTLA4 perform its conserved function (w= 0.42, <0.0005) through competing with CD28 on binding to CD80/CD86. The oldest component of this pathway is CD86 that first appeared in Arthropoda. Whereas this pathway is suggested to be depend on LFA1 which first appeared in Tunicate, CTLA4 has the ability to suppress conventional T cells through increasing the expression of IDO which first appeared in sponges or reducing the GCLC and GSS which also first appeared in sponges. Taken together these observations suggest that Tregs employed CTLA4 as soon as it appeared in vertebrates to suppress ancient pathways known to be responsible for cell survival. Interesting Tregs seems have adopted the conserved the ancient function of the CD40/CD40L pathways, but considerably adapted the ancient A20, neuropilin-1. Furthermore it used vertebrates LAG3 to suppress TGFB is a known suppressor that was shown to suppress [30]. It has diverged in sponges and evolved under a rapid positive evolution (w 1.46, <0.0001). Thus it seems that Tregs adapted the usage of Tregs for their own environments. Interestingly in the induction of apoptosis pathways, Tregs seemed to have a vertebrates innovation pattern except for usage of TRAIL. The pattern of regulating more ancient systems is also apparent in usage of FOXP3 which appeared after the emergence of Tbet. Conversely in the NFAT pathway, Tregs seems to have acquired an ancient pathways of ICER. This is also evident in calcium regulations pathways.

### Origin of Tregs suppressors

Our investigation supports the notion of the divergent origin of regulation of the adaptive immune system. We have found that CTLA4, LAG3 seem to have diverged from Ig-like proteins. On the other hand, TRAIL and CD40L seem to belong have diverged from tumor necrosis factor ligand superfamily member 10-like. Interestingly, GITR and CD40R diverged from Tumor Necrosis Factor Receptor Superfamily, highlighting the role of the TNF family and its receptors in the regulation of the adaptive immune system. This is further supported by the emergence of A20 from a Tnf aloha-induced family in sponges. However other origins also seem to have contributed d to Tregs suppression, such as Cub containing domain, and Trypsin (99%). Origin of regulation is also not restricted in vertebrates as several ancient origins are suggested for CD39 and CD73. One of the strongest examples is Calcineurin with origin suggested to exist in Hesseltinella vesiculosa fungi and Dictyostelium discoideum amoeba 4e-06. Take together our results suggest that the nature of regulation could have first appeared in the earliest metazoan and with various divergent mechanisms appearing at the same time.

Lampreys suppression of immune cells functions is intriguing. Lampreys has two main adaptive immune cells types that are analogues to T cells. In vertebrates, CTLA4 consist of five main components, where CTAL4 interact with its ligand CD28 along with CD80 and CD86. LFA1 also was shown to play an important role in Tregs CTLA4 mediated suppression as it is (does what?). A similar pathway to CTLA4 mediated suppression by Tregs vertebrates, is unlikely to occur in lampreys as it lacks CTAL4 as well as CD80, CD86 and CD28 as well as LFA1 (figure 1)[26].We found that TGFB2 is expressed in lampreys along with GP130 and TRAIL. Interestingly, FoxP3 does not seem to appear in lampreys suggesting that lampreys lack a FoxP3+ -LIKE Tregs. However, they possess an ICER proetin that is capable of regulating NFAT pathway. They also possess an Calcineurin (PPP3CA) that is capable of regulating Nfkb. Taken together, our results suggest that Lamprey posses a Treg-like cell, however its function is not mediated by FOXP3.

